# Context-dependent tonic signaling shapes the performance and manufacturability of a 4-1BB–based HER2 CAR-T cell therapy

**DOI:** 10.64898/2026.05.11.724226

**Authors:** Laura Angelats, Berta Marzal, Alba Rodriguez-Garcia, Marta Español-Rego, Teresa Lobo-Jarne, Maria Hernandez-Sanchez, Guim Cascalló, Salut Colell, Marta Giménez-Alejandre, Guillem Colell, Joan Castellsagué, Irene Andreu-Saumell, Hugo Calderón, Patricia Galván, Alvaro Urbano-Ispizua, Julio Delgado, Europa Azucena González-Navarro, Aleix Prat, Manel Juan, Sonia Guedan

## Abstract

The development of clinically effective CAR-T cell therapies for solid tumors requires careful optimization of receptor design, functional fitness, and manufacturability. While advancing low-affinity HER2-targeting CAR-T cells toward clinical application, we found that the candidate with the strongest in vivo antitumor activity—comprising a CD8α hinge and transmembrane region and a 4-1BB co-stimulatory domain—exhibited measurable tonic signaling. This basal antigen-independent signaling, likely driven by high CAR surface expression, was associated with increased apoptosis and reduced ex vivo expansion under research-grade manufacturing conditions. Modification of the transmembrane domain reduced CAR surface expression but did not alleviate tonic signaling and instead impaired antitumor activity. By contrast, transient pharmacologic inhibition of CAR signaling with dasatinib rescued expansion and reduced apoptosis in small-scale research cultures. Notably, these tonic-signaling-associated defects were largely absent during large-scale, GMP-compliant manufacturing, which enabled robust CAR-T cell expansion without additional benefit from dasatinib supplementation. Together, these findings show that tonic signaling is not inherently detrimental to CAR-T cell performance and that its functional consequences are highly dependent on manufacturing context. Our study underscores the importance of evaluating CAR candidates within clinically relevant production platforms and supports the advancement of this 4-1BB–based HER2-specific CAR-T cell product toward clinical testing.

## INTRODUCTION

Chimeric Antigen Receptor (CAR) T-cell therapy has emerged as a transformative modality in cancer immunotherapy, achieving remarkable clinical responses across several hematologic malignancies. Nevertheless, translating this success to solid tumors and achieving durable responses remain significant challenges[1, 2].

Effective CAR-T cell activity in solid tumors requires robust in vivo expansion and sustained persistence while avoiding premature T-cell exhaustion. Among the factors that can undermine CAR-T performance is tonic signaling, defined as antigen-independent, constitutive activation of CAR-T cells. Tonic signaling has been shown to drive dysfunctional T-cell states characterized by impaired proliferation, increased apoptosis, metabolic dysregulation, and accelerated differentiation toward exhaustion, ultimately limiting antitumor activity and persistence [3–5]. A central mechanism underlying tonic signaling is the self-aggregation and clustering of CAR molecules, which is strongly influenced by high CAR expression levels on the cell surface or scFv instability [3–7]. Structural features such as the length of the linker connecting the variable heavy and light chains of the scFv [8] and the architecture of the hinge region [9] can modulate CAR surface density or the tendency of CARs to cluster and signal autonomously. In addition, the choice of co-stimulatory domain has a major role in shaping both the phenotype and the intensity of tonic signaling. CD28-based CARs can drive antigen-independent proliferation and sustained cytokine secretion, promoting accelerated differentiation and exhaustion, resulting in reduced antitumor activity [4, 5, 10]. By contrast, tonic signaling in 4-1BB-containing CARs, particularly when overexpressed, can lead to increased T-cell apoptosis, thereby limiting T cell expansion and impairing antitumor activity [3, 11]. Several approaches have been developed to mitigate tonic signaling, including reducing CAR expression levels [3, 4, 12], enhancing scFv stability [7], tuning CAR charge density to prevent autonomous oligomerization [6], or pharmacologic modulation of CAR signaling [13–20]. However, although these studies often suggest that CARs prone to tonic signaling should be avoided, emerging evidence suggests that moderate levels of tonic signaling, particularly in 4-1BB-based CARs, may actually enhance T cell priming and support antitumor function [8]. These observations suggest that the biological consequences of tonic signaling are highly context dependent and may vary according to CAR architecture and manufacturing conditions. Importantly, although tonic signaling has been extensively studied in small-scale research settings, whether these observations translate to clinically relevant GMP-compliant manufacturing platforms remains poorly understood. Differences in T-cell activation, transduction, and expansion conditions during GMP manufacturing may substantially influence CAR expression levels, activation state, and the functional consequences of tonic signaling.

In this study, we sought to rationally optimize a low-affinity HER2-targeting CAR by dissecting how co-stimulatory domain selection and transmembrane configuration influence tonic signaling, expansion, and antitumor activity under both research-grade and clinical-grade GMP manufacturing conditions. We identified 4-1BB as the optimal co-stimulatory domain based on its superior in vivo efficacy and persistence, despite reduced ex vivo expansion associated with tonic-signaling-induced apoptosis. Modifying the CD8α-based transmembrane domain did not attenuate tonic signaling but instead impaired antitumor efficacy. By contrast, transient pharmacologic inhibition of CAR signaling with dasatinib effectively mitigated tonic signaling in research-grade cultures. Notably, the impact of tonic signaling was minimal during large-scale, GMP-compliant CAR-T manufacturing, eliminating the need for dasatinib supplementation in this setting. Together, these findings provide mechanistic insights and practical guidance for advancing 4-1BB–based HER2-specific CAR-T cells toward clinical translation and supported the selection of a lead CAR construct for further development. This construct, designated ARI-HER2, is planned for evaluation in a future phase 1 clinical trial.

## RESULTS

### 4-1BB co-stimulation enhances HER2 CAR-T cell efficacy despite impaired ex vivo expansion

To identify the optimal intracellular signaling domain for our HER2-specific CAR, we generated CAR constructs incorporating low affinity 4D5-5 scFv together with either 4-1BB, CD28, or ICOS co-stimulatory domains, all coupled to the CD3ζ intracellular signaling domain (**Figure 1A**). We also evaluated a mutant version of the CD28 co-stimulatory domain, CD28YMFM, previously reported to improve persistence in a different setting, was also tested [21].

**Figure 1.**
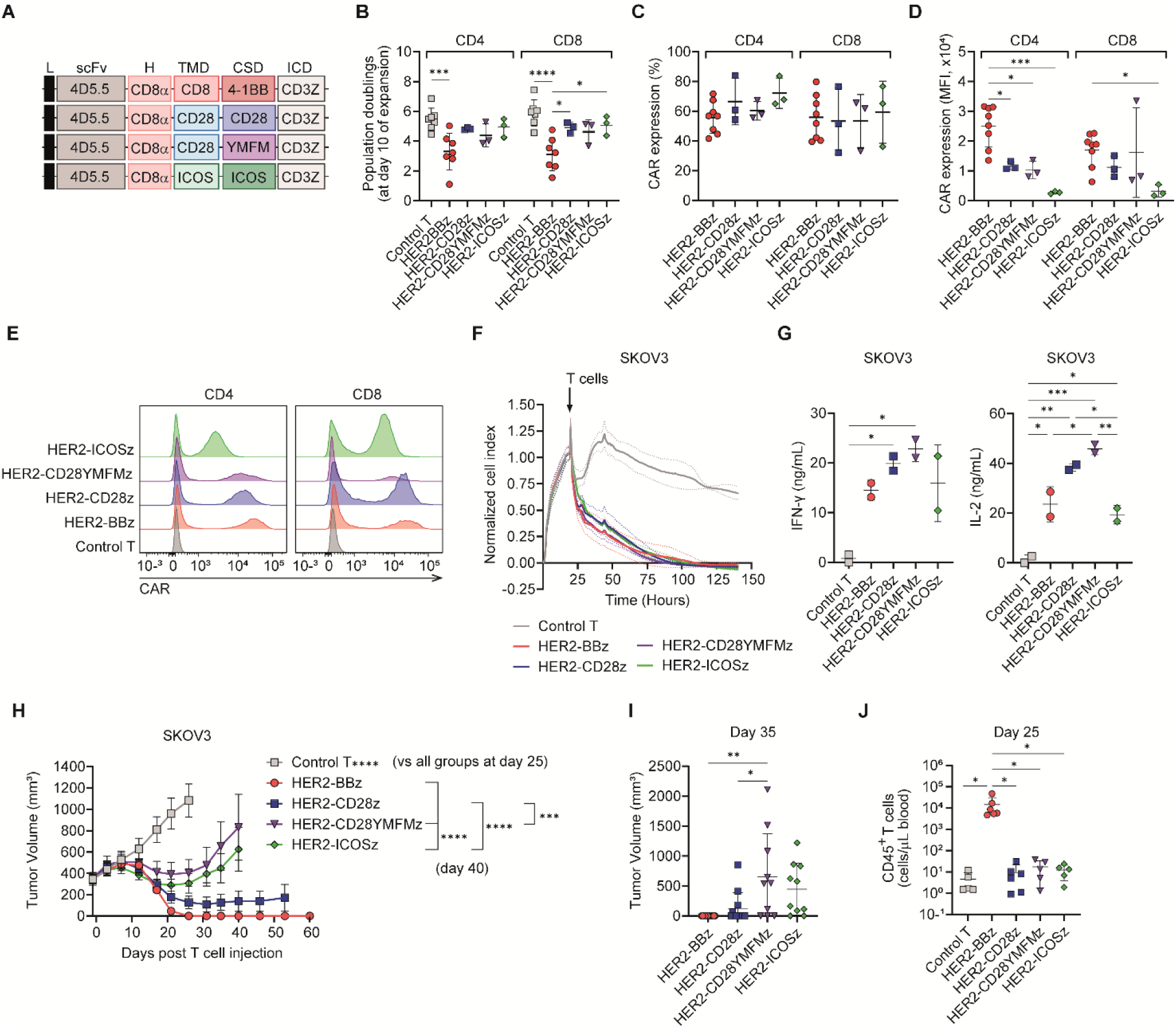
Functional comparison of HER2 CAR-T cells with distinct co-stimulatory domains identifies 4-1BB as the most effective despite reduced expansion. **(A)** Schematic of CAR constructs used. **(B)** Population doublings of CD4⁺ and CD8⁺ CAR-T cells at day 10 of expansion. Data are shown as mean ± SD and each dot represents a healthy donor (n=3-7). **(C)** Percentage and **(D)** mean fluorescence intensity (MFI) of CAR expression in CD4⁺ and CD8⁺ CAR-T cells at day 8 of expansion. Data are shown as mean ± SD and each dot represents a healthy donor (n=3-8). **(E)** Representative histogram of CAR expression. **(F)** Real-time cytotoxicity assay (xCELLigence) of CAR-T cells co-cultured with HER2⁺ SKOV3 ovarian cancer cells (effector-to-target ratio, E:T=1:1). The timepoint of T cell addition is indicated. Data are plotted as mean ± SD of normalized cell index (n=2 healthy donors). **(G)** IFN-γ (left panel) or IL-2 (right panel) levels after 24 h of co-culture of control T cells or indicated HER2 CAR-T cells with SKOV3 cells (E:T=3:1). Data are shown as mean ± SD and each dot represents a healthy donor (n=2). (H–J) *In vivo* evaluation in NSG mice bearing subcutaneous SKOV3 tumors treated with a single intravenous dose of 2×10⁶ CAR⁺ or control T cells. **(H)** Tumor growth kinetics following treatment. Data are shown as mean tumor over time volume ± SEM (n=10-12 tumors per group). **(I)** Tumor volumes at day 35 post-treatment. Data are shown as mean ± SD and each dot represents an individual tumor (n=10-12 tumors per group). **(J)** Persistence of CAR-T cells in peripheral blood at day 25 post-infusion. Data are shown as mean ± SD and each dot represents an individual mouse (n=5-6 mice per group). In **B**-**D** and **H**: *p<0.05, ***p<0.001, ****p<0.0001 by two-way ANOVA with Tukey’s multiple comparison test. In **G** and **I**-**J**: *p<0.05, **p<0.01, ***p<0.001 by one-way ANOVA with Tukey’s multiple comparison test. Abbreviations: L, leader peptide; H, hinge; TMD, transmembrane domain; CSD, costimulation domain; ICD, intracellular domain.

During *in vitro* expansion, CAR-T cells comprising the 4-1BB co-stimulatory domain exhibited significantly reduced population doublings compared to control T cells and CARs incorporating CD28, CD28YMFM or ICOS domains (**Figure 1B**). Although the frequency of CAR-expressing T cells was comparable across groups (**Figure 1C**), 4-1BB-based CAR-T cells displayed markedly higher CAR surface expression, as measured by mean fluorescence intensity (MFI) (**Figure 1D**-**E**). These findings suggested that elevated CAR surface density may promote tonic signaling and limit ex vivo expansion of 4-1BB-based HER2 CAR-T cells.

Next, we characterized the *in vitro* functionality of CAR-T cells following co-culture with HER2-expressing SKOV3 ovarian cancer cells. All groups exhibited comparable robust tumor cell killing (**Figure 1F**), and all CAR-T groups produced IFN-γ and IL-2 in co-culture, with CD28-and CD28YMFM-co-stimulated CAR-T cells showing the highest levels of cytokine secretion (**Figure 1G**). *In vivo* evaluation of antitumor efficacy was conducted in NSG mice bearing SKOV3 xenograft tumors. While all CAR constructs delayed tumor growth compared to control T cells, CARs containing CD28 or 4-1BB co-stimulatory domains showed the most substantial antitumor effects, with the HER2-BBz CAR achieving complete tumor eradication in all treated mice (**Figure 1H**-**I**). This superior antitumor response was correlated with a significant increase in T cell persistence (**Figure 1J**).

Together, these findings demonstrate that despite impaired ex vivo expansion, HER2-BBz CAR-T cells retain potent antitumor function and exhibit superior in vivo efficacy and persistence. Based on these results, we selected the 4-1BB co-stimulatory domain for further translational development of our HER2-specific CAR-T cell platform.

### Reducing CAR surface expression through transmembrane modification compromises in vivo efficacy

After identifying 4-1BB as the optimal intracellular co-stimulatory domain, we next investigated whether modification of the transmembrane (TM) domain could improve the expansion of HER2-BBz CAR-T cells by reducing tonic signaling. Because the TM domain can influence CAR surface expression and receptor clustering, we hypothesized that altering this region might attenuate tonic signaling and enhance ex vivo expansion. To test this, we generated HER2 CAR constructs incorporating the 4-1BB co-stimulatory domain together with TM domains derived from CD8α, CD28, or ICOS (**Figure 2A**).

**Figure 2.**
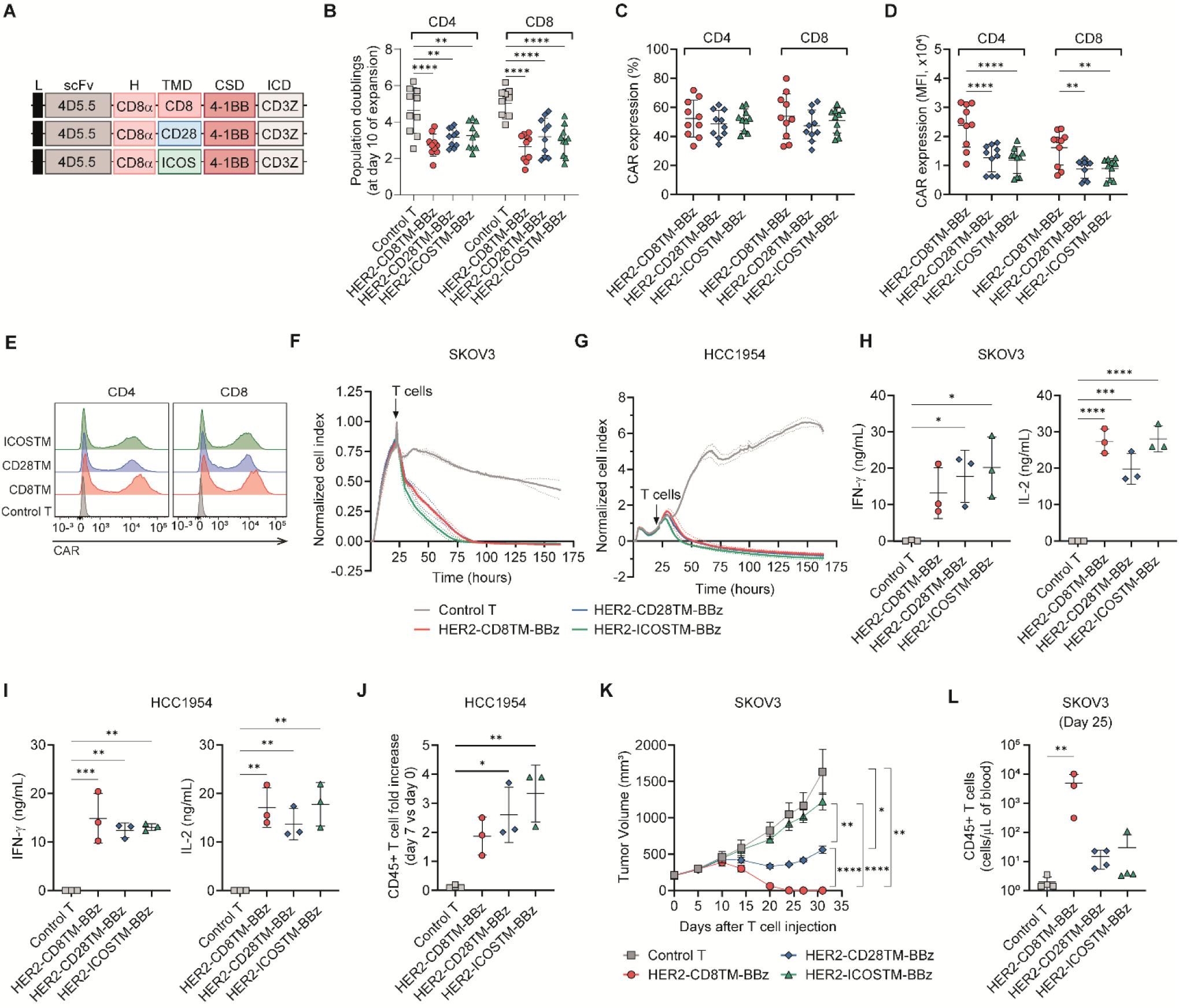
CD8α transmembrane domain substitution does not mitigate tonic signaling and compromises in vivo antitumor function of HER2-BBZ CAR-T Cells. **(A)** Schematic of CAR constructs used. **(B)** Population doublings of CD4⁺ and CD8⁺ CAR-T cells at day 10 of expansion. Data are shown as mean ± SD and each dot represents a healthy donor (n=10). **(C)** Percentage and **(D)** mean fluorescence intensity (MFI) of CAR expression in CD4⁺ and CD8⁺ CAR-T cells at day 8 of expansion. Data are shown as mean ± SD and each dot represents a healthy donor (n=10). **(E)** Representative histogram of CAR expression. **(F-G)** Real-time cytotoxicity assay (xCELLigence) of CAR-T cells co-cultured with HER2⁺ **(F)** SKOV3 ovarian or **(G)** HCC1954 breast cancer cells (E:T=1:1). The timepoint of T cell addition is indicated. Data are plotted as mean ± SD of normalized cell index (n=2 healthy donors). **(H-I)** IFN-γ (left panel) or IL-2 (right panel) levels after 24h of co-culture of control T cells or indicated HER2 CAR-T cells with **(H)** SKOV3 or **(I)** HCC1954 cells (E:T=3:1). Data are shown as mean ± SD and each dot represents a healthy donor (n=3). **(J)** T cell proliferation at day 7 after co-culture with HCC1954 tumor cells relative to day 0, as analysed by flow cytometry. Data are shown as mean ± SD and each dot represents a healthy donor (n=3). (K-L) *In vivo* evaluation in NSG mice bearing subcutaneous SKOV3 tumors treated with a single intravenous dose of 2×10⁶ CAR⁺ or control T cells. **(K)** Tumor growth kinetics following treatment. Data are shown as mean tumor volume ± SEM over time (n=6-8 tumors per group). **(L)** Persistence of CAR-T cells in peripheral blood at day 25 post-infusion. Data are shown as mean ± SD and each dot represents an individual mouse (n=3-4 mice per group). In **B**-**D** and **J**: *p<0.05, **p<0.01, ****p<0.0001 by two-way ANOVA with Tukey’s multiple comparison test. In **H**-**I** and **L**: *p<0.05, **p<0.01, ***p<0.001, ****p<0.0001 by one-way ANOVA with Tukey’s multiple comparison test. In **I**: *p<0.05, ** p<0.01 by one-way ANOVA with Holm-Sidak’s multiple comparison test. Abbreviations: L, leader peptide; H, hinge; TMD, transmembrane domain; CSD, costimulation domain; ICD, intracellular domain.

Although replacement of the CD8α TM with CD28 or ICOS TM domains showed a trend toward increased population doublings during ex vivo expansion, these differences did not reach statistical significance (**Figure 2B**). Notably, while different transmembrane domains did not affect the frequency of CAR^+^-T cells (**Figure 2C**), there were marked differences in expression intensity. Specifically, constructs with CD28 or ICOS TMs exhibited significantly reduced MFI compared to that with CD8α TM (**Figure 2D-E**).

We next characterized the CAR constructs *in vitro* following co-culture with ovarian cancer (SKOV3) and HER2+ breast cancer (HCC1954) cell lines. No significant differences between the CARs containing different transmembrane domains were observed in terms of cytotoxicity (**Figure 2F-G**) or cytokine secretion (e.g., IFN-γ and IL-2) (**Figure 2H-I**). However, proliferation assays upon antigen encounter revealed higher proliferation rates for CARs containing CD28 or ICOS TMs as compared to CD8α TM (**Figure 2J**).

We next analyzed the antitumor efficacy of 4-1BB-based CAR-T cells containing various transmembrane (TM) domains in NSG mice bearing SKOV3 tumors. Surprisingly, the transmembrane domain of the CAR played a crucial role in *in vivo* antitumor efficacy. Contrary to the *in vitro* proliferation data, CAR-T cells with an ICOS TM exhibited severely impaired antitumor efficacy, those with a CD28 TM showed only moderate efficacy, and CAR-T cells incorporating a CD8α TM were the only configuration that achieved complete tumor eradication in all treated mice (**Figure 2K**). This enhanced antitumor effect correlated with a significant increase in T cell persistence in the blood 25 days post-treatment (**Figure 2L**). Based on these findings, we selected the HER2-CD8α TM-BBz CAR for further preclinical characterization.

### Transient dasatinib treatment during ex-vivo expansion mitigates tonic signaling across 4-1BB-based CAR-T cells

After selecting the optimal CAR construct, we next sought to determine whether transient pharmacologic inhibition of CAR signaling could mitigate tonic signaling and improve the impaired expansion of HER2-BBz CAR-T cells. Previous studies have shown that tonic signaling in 4-1BB-based CAR-T cells can promote early apoptosis, progressive loss of CAR-expressing cells, and reduced ex vivo expansion[3]. We therefore hypothesized that transient inhibition of CAR signaling during the early phase of expansion might alleviate these effects and improve CAR-T cell fitness.

To test this, we treated HER2-BBz CAR-T cells with dasatinib, a Src-family kinase inhibitor reported to transiently suppress CAR signaling. Because apoptosis typically occurs at early time points following CAR expression, dasatinib was added only on days 3–4 of T cell expansion to minimize exposure while targeting the window when tonic signaling contributes most to cell death. Cells were treated with a range of concentrations from 6 to 100 nM to determine the minimal dose required to inhibit tonic signaling.

Dasatinib increased population doublings in a dose-dependent manner, with 50 nM representing the lowest concentration that significantly enhanced expansion compared with untreated CAR-T cells (**Figure 3A**). As previously suggested [3], we observed a decline in the frequency of HER2-BBz CAR-expressing cells from day 5 to day 11 (**Figure 3B**), while CAR density was similar between groups at the end of the expansion (**Figure 3C**). Dasatinib partially mitigated the decline in CAR⁺ frequency, as a significant increase in the percentage of CAR^+^-T cells at day 11 was observed starting from 12.5 nM of dasatinib as compared to CAR-T cells expanded without the drug (**Figure 3B**). In line with this, the decline in CAR^+^ cells between day 5 and day 11 was reduced, decreasing from a 1.85-fold reduction in untreated cultures to a 1.40-fold reduction at 100 nM dasatinib (**Figure 3D**). To determine whether this reduction in CAR frequency reflected selective outgrowth of untransduced cells or CAR downregulation, we quantified vector copy number per genome in sorted surface CAR⁺ and CAR⁻ subsets. CAR⁻ cells lacked vector copies across conditions, indicating that the decline in CAR⁺ frequency resulted from untransduced cell enrichment rather than CAR downregulation (**Figure 3E**). This pattern suggests that CAR-expressing cells, particularly those with high CAR density, may undergo apoptosis early during expansion [3]. Consistent with this, apoptosis analysis on day 6 revealed that T cells cultured without dasatinib exhibited elevated apoptosis, whereas dasatinib at ≥ 25 nM significantly reduced apoptosis to levels comparable to control T cells (**Figure 3F**). Based on these findings, we selected 50 nM dasatinib as the optimal concentration to enhance T cell expansion, preserve CAR expression and reduce apoptosis.

**Figure 3.**
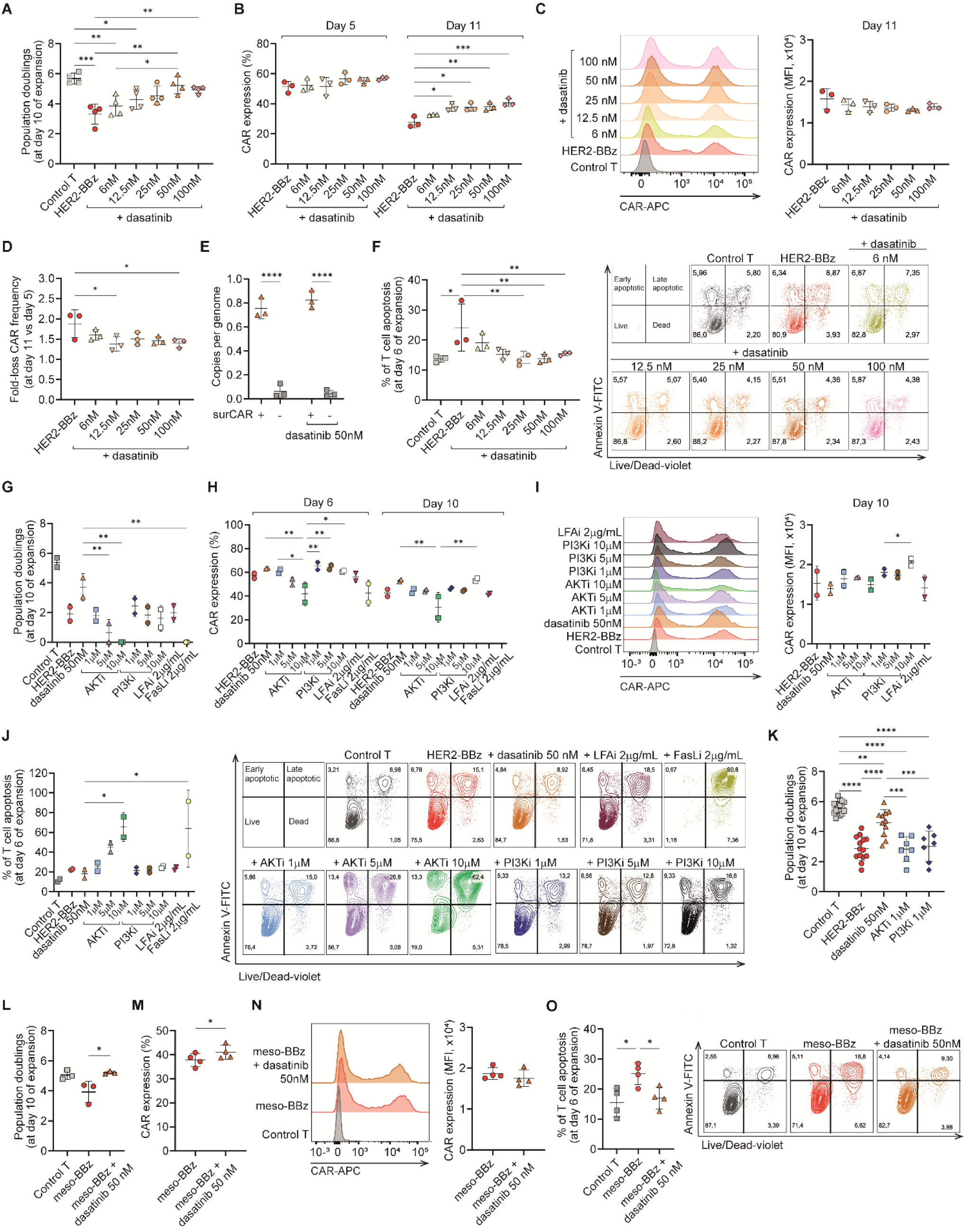
Transient dasatinib treatment during early expansion reduces tonic signaling and enhances 4-1BB CAR-T cell fitness. (A-F) HER2-BBz CAR-T cells were expanded in the absence or presence of increasing doses of dasatinib added to T cell cultures on days 3 and 4. **(A)** Population doublings of T cells at day 10 of expansion. Data are shown as mean ± SD, and each dot represents a healthy donor (n=4). **(B)** Percentage of CAR expression in T cells at days 5 and 11 of expansion. Data are shown as mean ± SD, and each dot represents a healthy donor (n=3). **(C)** Mean fluorescence intensity (MFI) of CAR expression at day 11 of expansion. A representative histogram is shown in the left panel. Data in the right panel are presented as mean ± SD, with each dot representing an individual healthy donor (n=3). **(D)** Fold-loss in CAR frequency at day 11 versus day 5. Data are shown as mean ± SD, and each dot represents a healthy donor (n=3). **(E)** Vector copy number per genome in HER2-BBz CAR-T cells cultured in the absence or presence of dasatinib (50 nM) and sorted for surface CAR expression into CAR^+^ and CAR^-^ populations, analysed at day 11 of expansion. Data represents mean ± SD and each dot represents a healthy donor (n=3). **(F)** Apoptotic T cells at day 6 of expansion as assessed by flow cytometry using Annexin V and live/dead staining. Apoptosis was defined as the sum of Dead/Annexin V⁺ cells. Data are shown as mean ± SD, and each dot represents a healthy donor (n=3). Representative flow cytometry plots are shown in the right panel. **(G-J)** HER2-BBz CAR-T cells were expanded in the absence or presence of the indicated inhibitors at the indicated concentrations, added on days 3 and 4. **(G)** Population doublings of T cells at day 10 of expansion. Data are shown as mean ± SD, and each dot represents a healthy donor (n=2). **(H)** Percentage of CAR expression in T cells at days 6 and 10 of expansion. Data are shown as mean ± SD, and each dot represents a healthy donor (n=2). **(I)** Mean fluorescence intensity (MFI) of CAR expression at day 10 of expansion. A representative histogram is shown in the left panel. Data in the right panel are presented as mean ± SD, with each dot representing an individual healthy donor (n=2). **(J)** Frequency of apoptotic T cells at day 6 of expansion, calculated as in (D). Data are shown as mean ± SD, and each dot represents a healthy donor (n=2). Representative flow cytometry plots are shown in the right panel. **(K)** Population doublings of HER2-BBz CAR-T cells expanded in the absence or presence of selected doses of the indicated inhibitors added on days 3 and 4, at day 10 of expansion. Data are shown as mean ± SD, and each dot represents a healthy donor (n=7-17). **(L-O)** Mesothelin-BBz CAR-T cells were expanded in the absence or presence of 50 nM dasatinib added on days 3 and 4. **(L)** Population doublings of T cells at day 10 of expansion. Data are shown as mean ± SD, and each dot represents a healthy donor (n=4). **(M)** Percentage of CAR expression in T cells at day 6 of expansion. Data are shown as mean ± SD, and each dot represents a healthy donor (n=4). **(N)** Mean fluorescence intensity (MFI) of CAR expression at day 10 of expansion. A representative histogram is shown in the left panel. Data in the right panel are presented as mean ± SD, with each dot representing an individual healthy donor (n=4). **(O)** Frequency of apoptotic T cells at day 6 of expansion, calculated as in (D). Data are shown as mean ± SD, and each dot represents a healthy donor (n=4). Representative flow cytometry plots are shown in the right panel. In **A**, **D, F-G**, **I**-**M**, and **O**: *p<0.05, **p<0.01, ***p<0.001, ****p<0.0001 by one-way ANOVA with Tukey’s multiple comparison test. In **B, E**, and **H**: *p<0.05, **p<0.01, ***p<0.001, ****p<0.0001 by two-way ANOVA with Tukey’s multiple comparison test.

We next compared dasatinib with additional pharmacological inhibitors previously reported to modulate CAR signaling [15–18]. Using the same treatment regimen (addition of drugs at day 3 and 4), we evaluated AKT and PI3K inhibitors across multiple concentrations during HER2-BBz CAR-T cell expansion. In parallel, compounds targeting LFA-1 and FasL were also tested. All inhibitors were directly benchmarked against dasatinib to assess their relative ability to support proliferation, maintain CAR expression, and limit apoptosis. None of the tested compounds significantly improved the expansion of HER2-BBz CAR-T cells, and dasatinib at 50 nM yielded the highest population doublings (**Figure 3G**). Regarding CAR expression, only PI3K inhibition at 10 μM retained CAR⁺ frequencies comparable to dasatinib (**Figure 3H**), and showed increased MFI at the end of the expansion (**Figure 3I**), but this dose showed trends towards reduced population doublings. No condition surpassed dasatinib in reducing apoptosis (**Figure 3J**). In fact, AKT inhibition at 5-10 μM and FasL inhibition impaired CAR-T cell expansion, decreased CAR expression, and increased apoptosis (**Figure 3G**-**3J**). To further confirm that dasatinib was the superior condition, we evaluated population doublings at the end of expansion in more than seven healthy donors using the selected concentrations of dasatinib (50 nM), AKTi (1 μM), and PI3Ki (1 μM). Only dasatinib significantly enhanced HER2-BBz CAR-T cell expansion across donors (**Figure 3K**).

Finally, to assess whether these effects were generalizable beyond the HER2-BBz construct, we evaluated the impact of dasatinib on 4-1BB-based CAR-T cells targeting mesothelin (M11). Transient exposure to 50 nM dasatinib during early expansion (days 3-4) significantly enhanced T-cell proliferation (**Figure 3L**), increased the proportion of CAR-expressing cells (**Figure 3M**) but similar expression levels (**Figure 3N**), and reduced apoptosis (**Figure 3O)** relative to untreated M11-BBz CAR-T cells. These findings indicate that the beneficial effects of transient dasatinib treatment are not restricted to a specific antigen-binding domain and may extend to additional CAR architectures.

### Transcient dasatinib exposure during *ex vivo* expansion of HER2-BBz CAR-T cells enhances cytokine production while preserving antitumor efficacy

We next sought to characterize the effector functions of HER2-BBz CAR-T cells expanded in the presence of dasatinib to ensure that exposure to increasing concentrations of the drug did not compromise their antitumoral activity. HER2-BBz CAR-T cells cultured with dasatinib displayed an increased production of IFN-γ upon co-culture with HER2-expressing SKOV3 ovarian cancer cells, with statistically significant differences detected starting at 25 nM (**Figure 4A**), compared with untreated CAR-T cells. Regarding cytotoxicity, dasatinib-treated HER2-BBz CAR-T cells lysed target cells as efficiently as untreated counterparts (**Figure 4B**). Consistent with these *in vitro* findings, *in vivo* antitumor efficacy was maintained in CAR-T cells expanded with dasatinib, even at high concentrations of 1000 nM, demonstrating that the drug did not impair their therapeutic potential (**Figure 4C**).

**Figure 4.**
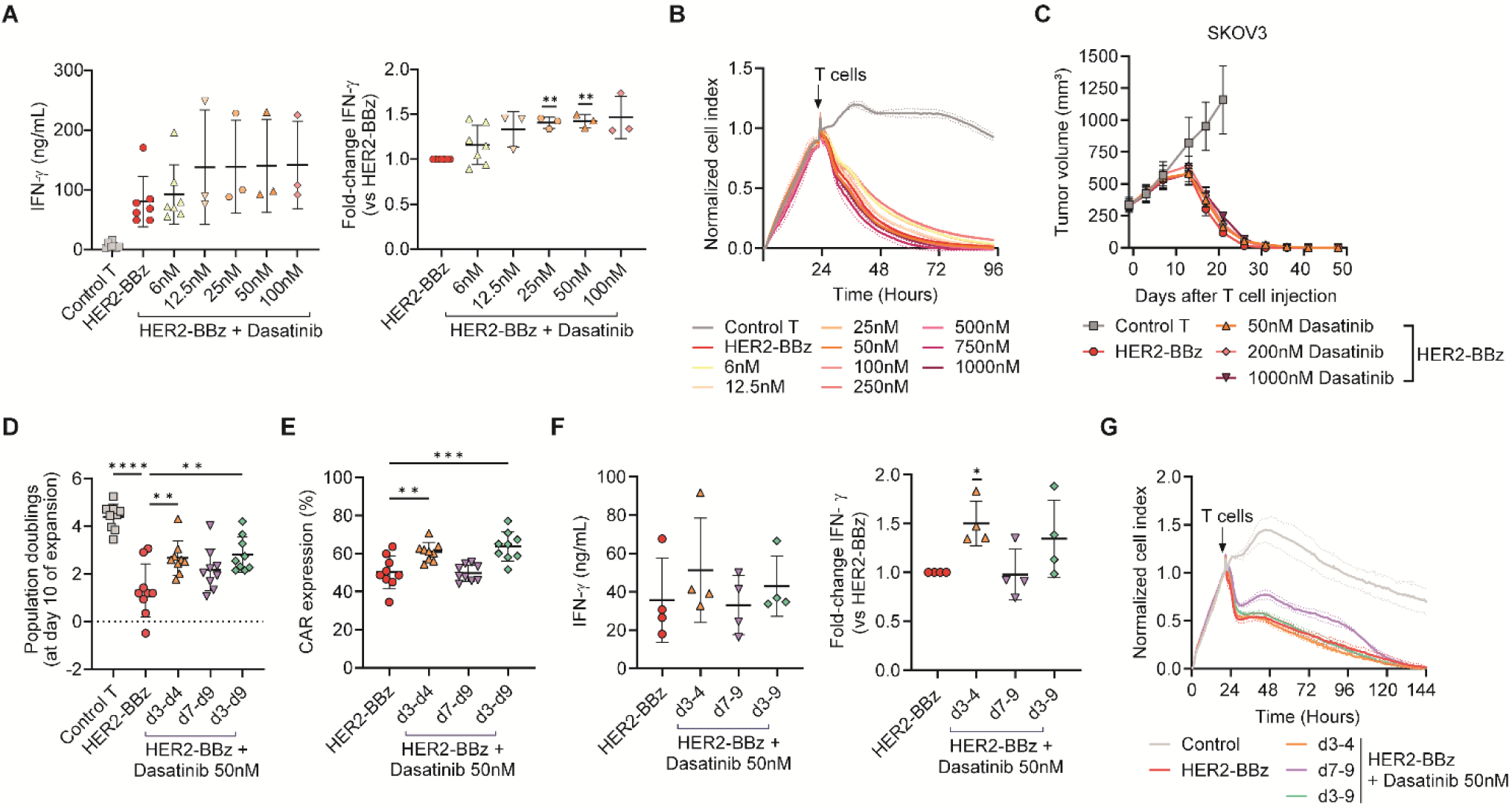
Early dasatinib treatment enhances cytokine production while preserving antitumor efficacy, whereas prolonged or delayed administration provides no additional benefit. (A-C) HER2-BBz CAR-T cells were expanded in the absence or presence of increasing doses of dasatinib added to T cell cultures on days 3 and 4. **(A)** IFN-γ production after 24h of co-culture with SKOV3 cells (E:T=3:1). Absolute IFN-γ levels (left) and fold change relative to untreated HER2-BBz CAR-T cells (right) are shown. Data represent mean ± SD, and each dot indicates an individual healthy donor (n=3-7). **(B)** Real-time cytotoxicity assay (xCELLigence) of CAR-T cells co-cultured with SKOV3 ovarian cancer cells (E:T=1:1). The timepoint of T cell addition is indicated. Data are shown as mean ± SD of the normalized cell index (n=2 healthy donors). (C) *In vivo* evaluation in NSG mice bearing subcutaneous SKOV3 tumors treated with a single intravenous dose of 2×10⁶ CAR⁺ or control T cells. Tumor growth kinetics are shown as mean tumor volume ± SEM over time (n=8 tumors per group). **(D**-**G)** HER2-BBz CAR-T cells were expanded in the presence of 50 nM dasatinib added at different time intervals (days 3–4, days 7–9, or days 3–9). **(D)** Population doublings at day 10 of expansion. Data represent mean ± SD, and each dot represents a healthy donor (n=9). **(E)** Percentage of CAR expression in T cells at day 8 of expansion. Data represent mean ± SD, and each dot represents a healthy donor (n=9). **(F)** IFN-γ production after 24h of co-culture with SKOV3 cells (E:T=3:1). Absolute IFN-γ levels (left) and fold change relative to untreated HER2-BBz CAR-T cells (right) are shown. Data represent mean ± SD and each dot represents a healthy donor (n=4). **(G)** Real-time cytotoxicity assay (xCELLigence) of CAR-T cells co-cultured with SKOV3 ovarian cancer cells (E:T=1:1). The timepoint of T cell addition is indicated. Data are plotted as mean ± SD of normalized cell index (n=2 healthy donors). In **A** and **F**: *p<0.05, **p<0.01 by one-sample t-test. In **D**-**E**: **p<0.01, ***p<0.001, ****p<0.0001 by one-way ANOVA with Tukey’s multiple comparison test.

### Early transient dasatinib exposure is sufficient to improve HER2-BBz CAR-T cell expansion

We then aimed at exploring whether altering the timing or duration of dasatinib exposure could differentially affect HER2-BBz CAR-T cell expansion. To do so, we compared the standard early exposure (days 3-4) with later exposure (days 7-9, similar to the regimen described by Zhang et al., [13]) and an extended continuous exposure from days 3 to 9. On day 10 of expansion, both early (d3-4) and extended (d3-9) dasatinib treatment significantly increased population doublings relative to untreated controls, whereas later exposure (d7-9) produced only a mild, non-significant effect (**Figure 4D**).

A similar pattern was observed for CAR expression: CAR⁺ frequencies at the end of expansion were significantly increased only in conditions where dasatinib was applied beginning on day 3 (**Figure 4E**). Likewise, enhanced IFN-γ secretion was maintained only when dasatinib was administered during the early expansion window, whereas treatment starting on day 7 failed to preserve this effect (**Figure 4F**). *In vitro* cytotoxicity against target cells remained comparable across all conditions (**Figure 4G**).

Together, these results indicate that early, transient inhibition of tonic CAR signaling is essential for improving HER2-BBz CAR-T cell expansion and maintaining CAR expression. In contrast, late or extended dasatinib exposure does not provide additional benefit and may be unnecessary. Based on these findings, treatment with 50 nM dasatinib on days 3-4 remains the optimal regimen.

### Dasatinib reduces apoptotic transcriptional signatures in HER2-BBz CAR-T cells

To uncover transcriptional changes underlying the effects of dasatinib during *ex vivo* expansion, we profiled gene expression in untransduced control T cells (UTD) and HER2-BBz CAR-T cells expanded with or without dasatinib using the nCounter® CAR-T Characterization Gene Expression Panel (NanoString Technologies). HER2-BBz CAR-T cells expanded without dasatinib exhibited the highest number of differentially expressed genes (122) compared to control T cells (**Figure 5A**). This transcriptional program was characterized by the significant upregulation of genes associated with T-cell activation and effector function (CD247, IL2RA, TNFRSF4, TNFRSF9, IFNG, GZMB) or interferon-stimulated genes (IRF1, IRF7, IFIT3, OAS1), together with increased expression of activation of NF-κΒ and AP-1 signaling pathways (NFKB2, NFKBIA, MAP3K14, FOS, JUN) and regulatory factors (CTLA4, TIGIT, FOXP3, IKZF2). In addition, stress-and apoptosis-related genes (DDIT4, XAF1, TNFRSF10B) were also upregulated. Together, these features are consistent with antigen-independent tonic signaling-induced activation. Dasatinib treatment diminished the magnitude of this activation program, evidenced by a reduction in the number of differentially expressed genes (68) compared to control T cells (**Figure 5B**). While CAR-T cells retained expression of activation and signaling-related genes (CD247, TNFRSF4, TNFRSF9, NFKB2, JUN), dasatinib-treated cells exhibited a shift toward a less inflammatory and less cytotoxic profile, including reduced representation of immediate early genes and effector molecules, such as FOS, FOSB, GZMB and IL2. Direct comparison between HER2-BBz CAR-T cells cultured in the presence or absence of dasatinib confirmed a significant downregulation of genes involved in activation (FOS, FOSB), effector function (GZMB, IL2), inflammatory signaling (CXCL8, CSF2) and stress/apoptosis pathways (DDIT4) (**Figure 5C**). To better understand these changes, we performed gene ontology enrichment analysis focusing on the apoptotic process pathway (GO:0006915). This pathway was significantly upregulated in untreated HER2-BBz CAR-T cells relative to control T cells (**Figure 5D**). In contrast, dasatinib-treated HER2-BBz CAR-T cells had their apoptotic pathway downregulated compared with their untreated counterparts (**Figure 5E**). Together, these findings support the conclusion that transient dasatinib treatment during early expansion attenuates tonic-signaling-associated transcriptional programs and reduces apoptosis-related signatures in 4-1BB–based CAR-T cells at the transcriptional level.

**Figure 5.**
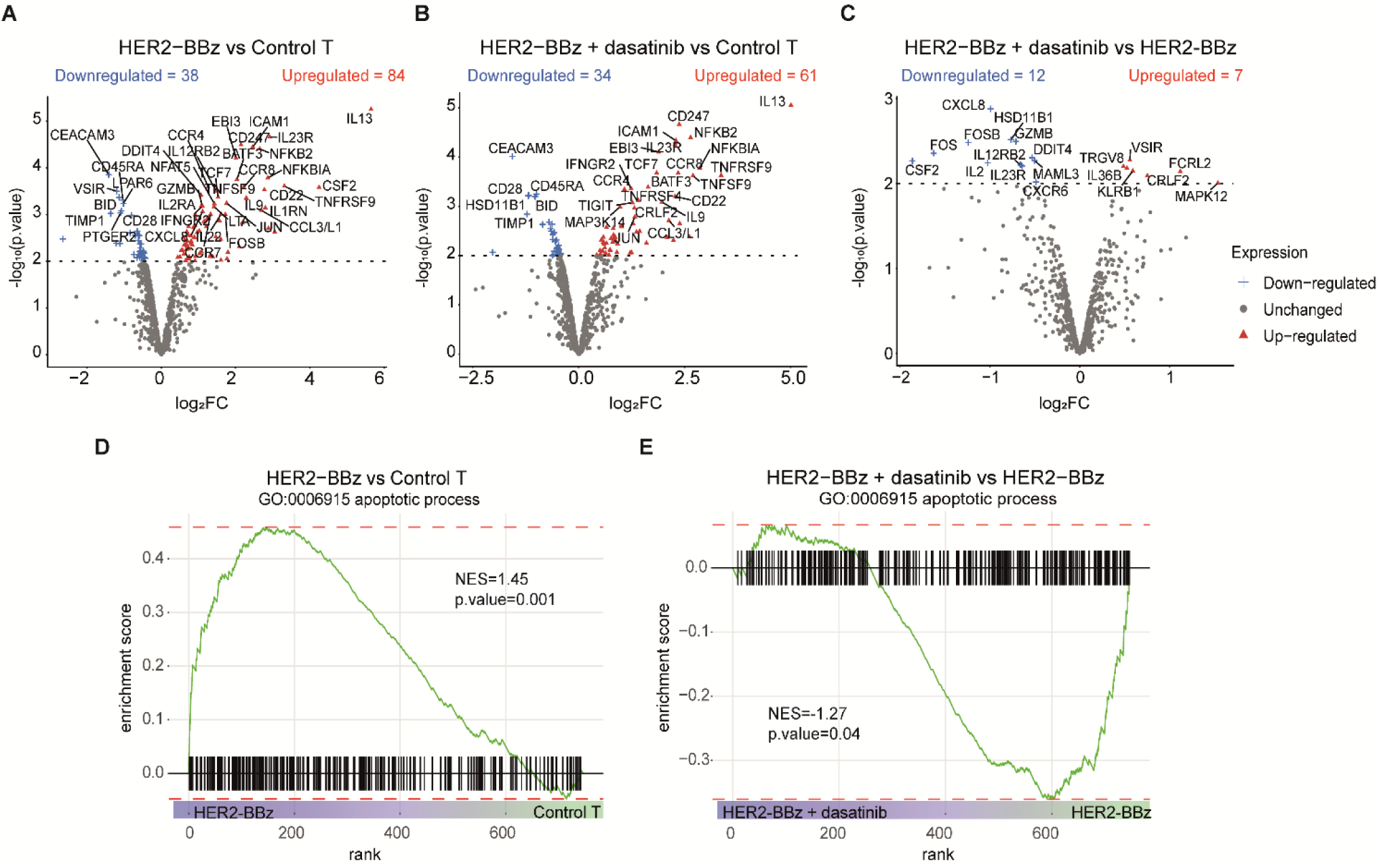
Dasatinib reduces apoptotic transcriptional signatures in HER2-BBz CAR-T cells. Gene expression profiling of untransduced control T cells and HER2-BBz CAR-T cells expanded with or without 50 nM dasatinib was performed using the nCounter® CAR-T Characterization Gene Expression Panel (NanoString Technologies) (n=2 paired-healthy donors per condition). **(A-C)** Volcano plots showing differential gene expression between **(A)** untreated HER2-BBz CAR-T cells and control T cells, **(B)** HER2-BBz CAR-T cells treated with dasatinib and control T cells, and **(C)** HER2-BBz CAR-T cells with versus without dasatinib. Upregulated genes are shown in red, downregulated genes in blue, and non-significant genes in grey. Selected significantly differentially expressed genes are labeled. The horizontal dotted line indicates a p-value threshold of 0.01. **(D-E)** Gene set enrichment analysis (GSEA) of the Gene Ontology apoptotic process pathway (GO:0006915) using significantly differentially expressed genes from **(D)** HER2-BBz CAR-T cells versus control T cells, and **(E)** HER2-BBz CAR-T cells with versus without dasatinib.

### Dasatinib supplementation is dispensable during GMP-compliant CAR-T cell manufacturing

We finally aimed to assess the impact of dasatinib supplementation on GMP-grade CAR-T cell production. For that, leukapheresis were split into paired cultures and processed on the CliniMACS Prodigy platform either with or without dasatinib (50 nM on days 3 and 5). CAR-T products were successfully manufactured in both conditions, and no significant differences were observed between dasatinib-treated and untreated products in terms of total cell numbers (**Figure 6A**) or CAR expression levels (**Figure 6B**) at the end of manufacturing. Additional product quality attributes including cell viability, CD3 purity, CD4:CD8 composition, and T cell phenotype, also remained comparable between treated and untreated products (**Figure 6C**-**G**). We next assessed functional potency by performing a cytotoxicity assay using HCC1954 target cells, showing similar killing capacity across conditions (**Figure 6H**). Consistently, Nanostring transcriptomic profiling identified no significant differences associated with dasatinib treatment (**Figure 6I**-**K**), indicating that the intervention did not induce major transcriptional changes during GMP processing.

**Figure 6.**
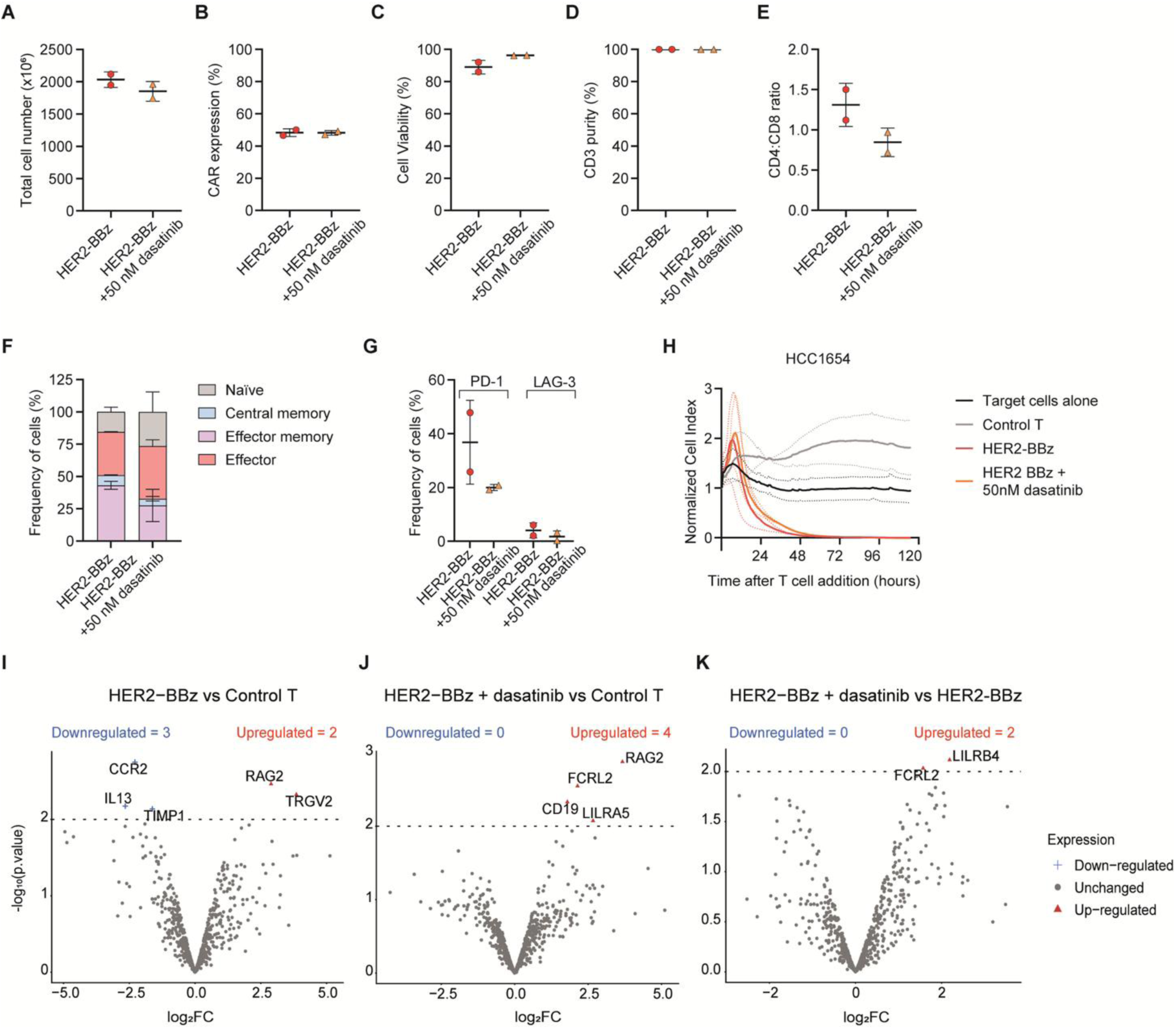
Dasatinib supplementation is not required for GMP-grade CAR-T cell manufacturing. Patient leukapheresis products were split into paired cultures and processed on the CliniMACS Prodigy platform either with or without dasatinib (50 nM on days 3 and 5). **(A)** Total cell yield, **(B)** percentage of CAR-expressing cells, **(C)** cell viability, **(D)** CD3^+^ T cell purity, and **(E)** CD4:CD8 ratio were assessed at the end of manufacturing. **(F)** CAR-T cell differentiation phenotype at the end of manufacturing, defined by CCR7 and CD45RA expression: naïve (CCR7^+^/CD45RA^+^), central memory (CCR7^+^/CD45RA^-^), effector memory (CCR7^-^/CD45RA^-^), and effector (CCR7^-^/CD45RA^+^). Frequencies of each subset are shown as mean ± SD (n=2). **(G)** Frequency of PD1^+^ or LAG3^+^ T-cells at the end of manufacturing. For panels **A**-**E** and **G**, data are presented as mean ± SD, with each dot representing an individual donor (n=2). **(H)** Real-time cytotoxicity assay (xCELLigence) of CAR-T cells co-cultured with HER2⁺ HCC1954 cancer cells (E:T=1:1). Data are plotted as mean ± SEM of normalized cell index (n=2 donors). **(I**-**K)** Gene expression profiling of untransduced control T cells and HER2-BBz CAR-T cells expanded with or without 50 nM dasatinib was performed using the nCounter® CAR-T Characterization Gene Expression Panel (NanoString Technologies) (n=2 paired donors per condition). Volcano plots showing differential gene expression between **(I)** untreated HER2-BBz CAR-T cells and control T cells, **(J)** HER2-BBz CAR-T cells treated with dasatinib and control T cells, and **(K)** HER2-BBz CAR-T cells with versus without dasatinib. Upregulated genes are shown in red, downregulated genes in blue, and non-significant genes in grey. Selected significantly differentially expressed genes are labeled. The horizontal dotted line indicates a p-value threshold of 0.01.

Taken together, these findings show that although dasatinib successfully mitigated tonic signaling in research-grade conditions, its addition did not enhance the quality, phenotype, or functional performance of HER2-BBZ CAR-T cell product during GMP-grade manufacturing, indicating that the baseline GMP process already produced high-quality cells without the need for dasatinib supplementation.

## DISCUSSION

In this study, we sought to develop a HER2-specific CAR-T therapy with potential for clinical translation. Among the CAR constructs evaluated, the combination of a CD8α hinge and transmembrane domain with a 4-1BB co-stimulatory domain conferred the most favorable antitumor properties. However, this configuration was also associated with reduced ex vivo expansion, likely as a consequence of tonic signaling.

Tonic signaling in CARs incorporating the 4-1BB costimulatory domain, has previously been linked to impaired ex vivo expansion, associated with high CAR surface density leading to increased apoptosis, and progressive downregulation of CAR expression [3]. Our findings are consistent with these observations, reinforcing the notion that excessive basal signaling can negatively impact T cell proliferative capacity during expansion. In spite of this, HER2-BBZ CAR-T cells exhibited superior in vivo performance, characterized by enhanced antitumor efficacy and long-term persistence. While this may seem paradoxical, accumulating evidence suggests that tonic signaling, when maintained within an optimal range, can be functionally advantageous. In particular, 4-1BB-mediated tonic signaling has been associated with improved T cell persistence and sustained antitumor activity [8, 22]. Together, these findings support the concept that the qualitative and quantitative features of tonic signaling, rather than its mere presence, are critical determinants of CAR-T cell fitness and therapeutic efficacy.

Because different co-stimulatory domains have different patterns of tonic signaling, we explored alternative configurations. Tonic signaling in CD28-based CARs has been associated with T cell dysfunction [4, 5, 10]. In our study, CD28-based HER2-specific CAR-T cells expanded efficiently ex vivo, expressed significantly lower CAR levels compared to 4-1BB-based constructs, and showed only a trend toward increased cytokine production. Despite this, reduced antitumor efficacy and decreased persistence in vivo was consistent with a dysfunctional phenotype, as previously reported by our group using this CAR construct [23]. Moreover, incorporation of the mutant CD28-YMFM or ICOS costimulatory domains failed to improve persistence or antitumor activity in our model, in contrast to prior studies demonstrating enhanced durability and tumor control [11, 21]. These discrepancies may be attributable to differences in CAR design, particularly scFv affinity. Previous studies were conducted using a high-affinity anti-mesothelin scFv (SS1), whereas our work employs a low-affinity anti-HER2 scFv (4D5.5). Consistent with this notion, Chen et al. reported that a HER2-CD28Z CAR incorporating the high-affinity 4D5 scFv displayed elevated tonic signaling and a high exhaustion index, comparable to a CAR based on the SS1 scFv, although a direct comparison with the low-affinity 4D5.5 variant used in our study was not included [6]. Lower-affinity CARs are expected to deliver reduced signaling strength, which may result in a milder exhaustion phenotype [24]. In this context, the functional advantage conferred by modified CD28 signaling or ICOS co-stimulation may be attenuated, potentially explaining the lack of improvement observed in our system.

As previously stated, the tonic signaling observed in our 4-1BB-based CAR-T cells is most likely driven by high CAR expression levels. Beyond promoting tonic signaling, high CAR density has also been associated with poorer clinical outcomes [25]. In this context, and considering potential manufacturing and scalability challenges with our HER2-BBZ CAR-T cells, we explored strategies to mitigate tonic signaling. Although relatively few studies have directly addressed the contribution of the transmembrane domain to tonic signaling, this region is known to play a critical role in regulating CAR surface expression and stability [26]. Indeed, reducing CAR density through transmembrane domain modification has been shown to attenuate activation and cytokine production, thereby limiting CAR-associated toxicities [27–29]. Given the link between high CAR surface expression, activation and tonic signaling, we hypothesized that altering the transmembrane domain could reduce CAR density and thereby attenuate tonic signaling. In our study, replacement of the CD8a transmembrane domain in 4-1BB-based CARs with domains derived from CD28 or ICOS effectively reduced CAR surface expression. However, this decrease translated only into a modest trend toward increased population doublings, suggesting that tonic signaling was not substantially alleviated. These findings indicate that CAR surface density alone is unlikely to be the sole driver of tonic signaling, and that additional structural or signaling features may contribute to this phenomenon. Importantly, while in vitro effector functions such as cytokine secretion remained intact, substitution of the transmembrane domain markedly impaired antitumor activity and persistence in vivo. Notably, the persistence advantage conferred by 4-1BB signaling was lost when CD28-or ICOS-derived transmembrane domains were used. Our observations are consistent with prior studies showing that optimal CAR performance depends on appropriate pairing of transmembrane and intracellular domains, as exemplified by ICOS-based CARs requiring an ICOS transmembrane domain to persistence and overall anti-tumor efficacy [11].

Emerging strategies that incorporate small molecule inhibitors of T cell signaling pathways during CAR-T production have been proposed as a means to enhance memory phenotype and limit exhaustion [13–17, 19, 20, 30]. Building on this concept, we evaluated whether early pharmacologic inhibition of T-cell signaling could mitigate tonic signaling during CAR-T cell expansion. Among all inhibitors tested, dasatinib at 50 nM emerged as the most effective, markedly improving T cell yield, reducing apoptosis, and preventing CAR-T cell loss throughout the expansion. This reversible inhibition provides a controlled window in which tonic signaling is suppressed, supporting efficient proliferation and robust expansion.

A striking finding of our study was the limited translatability of observations related to tonic signaling between small-scale, research-grade expansion conditions and large-scale, GMP-compliant manufacturing. Under laboratory conditions, HER2-BBZ CAR-T cells showed poor expansion and increased apoptosis and required dasatinib supplementation to achieve sufficient yields. In contrast, manufacturing using the CliniMACS Prodigy automated cell processing system enabled robust expansion without dasatinib, yielding products that met release criteria. These discrepancies likely reflect fundamental differences in manufacturing parameters, including T cell activation and transduction conditions [31, 32]. Notably, T cell activation is a key determinant of CAR-T cell differentiation, exhaustion, and fitness, and varies substantially between platforms. While CD3/CD28 magnetic beads were used in our research-scale protocol, GMP manufacturing relied on TransAct, a polymeric nanomatrix that may provide a milder and more controlled activation stimulus. In addition, differences in transduction conditions between protocols may result in variable CAR expression levels, which may further influence the extent and functional consequences of tonic signaling.

These findings reinforce the concept that CAR-T cell behavior is not determined solely by CAR design, but is also strongly influenced by the manufacturing context. Accordingly, our dasatinib-based strategy may be particularly useful for supporting the expansion and preclinical evaluation of CAR constructs prone to tonic signaling or fratricide under research-grade conditions. However, our results suggest that the impact of such interventions may be diminished, or potentially unnecessary, in optimized large-scale GMP-compliant manufacturing workflows. While several studies as mentioned above have demonstrated the benefits of signaling inhibition at the preclinical level, translation to clinical-scale manufacturing remains limited and often lacks direct comparisons with conventional processes. For instance, studies scaling AKT inhibition into GMP manufacturing have reported CAR-T products with improved phenotypic fitness and in vivo functionality, although accompanied by reduced ex vivo expansion, highlighting trade-offs associated with such approaches [31, 33]. Similarly, clinical studies incorporating PI3K pathway modulation have shown encouraging antitumor activity and memory-enriched phenotypes, but typically without direct side-by-side comparisons to non-modified manufacturing, making it difficult to isolate the specific contribution of the pharmacological intervention [34]. In the case of dasatinib, preclinical studies have demonstrated its ability to reversibly inhibit proximal CAR signaling and prevent or reverse tonic signaling–associated exhaustion. Dasatinib has been incorporated into CAR-T manufacturing in recent clinical trials reporting phenotypic composition of manufactured products showing a predominance of central memory T cells and promising antitumor activity in solid tumors [35, 36], although, again, direct comparisons with and without dasatinib are lacking, and the extent to which preclinical observations translate to clinical benefit remains to be fully established. Notably, the CAR used in these studies is based on the GD2-targeting 14G2a scFv, which has been associated with clustering and tonic signaling. In contrast, our HER2 CAR uses the low-affinity 4D5-5 scFv, with the same CD8α transmembrane and 4-1BB intracellular domains, differing primarily in the antigen-binding region. Given that GD2 scFvs are reported to have a higher propensity to aggregate, whereas the behavior of 4D5-5 is less well defined, such intrinsic scFv properties may contribute to variability in tonic signaling and in the response to interventions such as dasatinib.

Collectively, our findings suggest that tonic signaling is not solely an intrinsic property of CAR design, but rather emerges from the interplay between receptor architecture and manufacturing context, underscoring the need to evaluate novel CAR strategies within clinically relevant production platforms. Within this framework, our work informed the selection of a lead 4-1BB–based HER2 CAR construct for future clinical development, where it will be referred to as ARI-HER2.

## MATERIAL AND METHODS

### Study approval

All mouse studies were performed under a protocol (184-20) approved by the Ethic Committee for Animal Experimentation (CEEA) of the University of Barcelona and the Generalitat de Catalunya. Mice were bred and housed at the University of Barcelona’s Animal Facility.

### Cell line culture

Human tumor cell lines SKOV3 (ovarian cystadenocarcinoma) and HCC1954 (breast ductal carcinoma) were purchased from American Tissue Culture Collection (Manassas, Virginia, USA). The lentiviral packaging line HEK293FT (human embryonic kidney) was purchased from Thermo Fisher Scientific (Waltham, Massachusetts, United States). The human T-lymphocyte line Jurkat Clone E6-1 (acute T-cell leukemia) was obtained from Sigma-Aldrich (Burlington, Massachusetts, USA). SKOV3 and HEK293FT cells were cultured in DMEM (Gibco) and HCC1954 and Jurkat cells in RPMI-1640 (Gibco). All media were supplemented with 10% heat-inactivated FBS (Merck) and penicillin/streptomycin (100 U/mL and 100 µg/mL, 1%, Gibco). HEK293FT cultures were further supplemented with 1% GlutaMAX and 1% non-essential amino acids (NEAA, Gibco). All cell lines were cultured at 37 °C in 5% CO₂, routinely authenticated, and confirmed mycoplasma-free.

### CAR construction and lentiviral production

All CAR sequences were synthesized by Genscript and cloned into the third-generation lentiviral vector pCCL under the control of EF1α promoter [37]. The second-generation HER2-specific CAR incorporates a low-affinity humanized 4D5-5 single-chain variable fragment (scFv) [38] fused to a CD8α leader peptide and followed by CD8α-derived hinge and transmembrane domains, 4-1BB costimulatory domain and CD3ζ signaling domain. Additional CAR constructs incorporating transmembrane domains from CD28 or ICOS, and containing CD28, CD28 YMFM [21] or ICOS [39] replacing 4-1BB, were also used. The mesothelin-directed CAR incorporated the M11 scFv (extracted from patent WO2015090230A1260) linked to CD8α hinge and transmembrane domains, 4-1BB and CD3ζ intracellular domains.

Lentiviral vectors were produced by transfecting HEK293FT and titrated on Jurkat cells. Briefly, HEK293FT cells were seeded at 10×10^6^ in a total volume of 18 mL of medium in p150 culture plates. After 18 hours, cells were transfected with 18 μg of the pCCL transfer plasmid (containing the CAR construct) together with a premixed packaging mix containing 15 μg of pREV, 15 μg of pRRE and 7 μg of pVSV using PEI® (Polysciences). Viral supernatant were harvested at 48 and 72h post-transfection, filtered through 0.45µm filters, concentrated using Lenti-X Concentrator (Takara Bio) as per manufacturer’s protocol, titrated by limiting dilution in Jurkat cells based on CAR surface expression, and stored at −80°C until use.

### Isolation, transduction and CAR T cell expansion

Human T cells were isolated from healthy donor buffy coats obtained from Banc de Sang i Teixits (Barcelona) under protocols approved by institutional review boards. CAR-T cells were generated and expanded, as previously described [40]. Briefly, human T lymphocytes were isolated using Lymphoprep (StemCell Technologies). CD4⁺ and CD8⁺ subsets were negatively isolated using RosetteSepTM Human CD4+ or CD8+ T-Cell Enrichment Cocktails (Stem Cell Technologies) and activated with CD3/CD28 Dynabeads (Thermo Fisher Scientific) at a 1:1 bead-to-cell ratio for CD4⁺ T cells and 2:1 for CD8⁺ T cells in the presence of human IL-7 and IL-15 (Miltenyi biotec), each at 10 ng/mL. 24 hours after activation, T cells were transduced with CAR-encoding lentiviral vectors. Dynabeads were removed on day 3, and T cells were subsequently expanded for 10 days in complete RPMI containing 10% FBS, 1% penicillin/streptomycin, 10 mM HEPES, 10 mM GlutaMAX, and cytokines IL-7 and IL-15 (10 ng/mL each, Miltenyi Biotec). On day 10, CAR T cells were cryopreserved in freezing medium (50% X-VIVO 15, 40% FBS, 10% DMSO) at a CD4:CD8 ratio of 1:1.

For the indicated experimental conditions, cells were treated at the specified time points with the following inhibitors: AKT inhibitor (Merck) at 1, 5, or 10 µM; PI3K inhibitor (Selleck Chemicals) at 1, 5, or 10 µM; LFA inhibitor (BioXCell) at 2 µg/mL; Fas ligand inhibitor (BD Biosciences) at 2 µg/mL; or dasatinib (BMS) at concentrations ranging from 6 to 1000 nM.

### Flow cytometry and cell sorting

For surface CAR detection, T cells were first labeled with the fixable viability dye eFluor 450 (eBioscience, Thermo Fisher) to exclude dead cells. Staining was performed in FACS buffer (2% FBS in PBS). Cells were incubated for 30 min at 4 °C in the dark with biotin-SP-AffiniPure F(ab′)₂ fragment-specific goat anti-human IgG (Jackson ImmunoResearch). After washing, cells were stained with streptavidin-PE or streptavidin-APC (BD Biosciences) together with fluorochrome-conjugated antibodies: CD4-BV510 or CD8-APC (BioLegend), CD4-FITC, CD8-APC-H7 or CD45-PerCP-Cy5.5 (BD Biosciences). At indicated studies, CAR^+^ and CAR^-^ T cell populations were separated using FACSAriaII or FACS Aria SORP (BD) cell sorters.

For apoptosis analysis, cells were first stained for surface CAR as described above. At the end of this procedure, samples were washed once with PBS, and then once with 1× binding buffer (Thermo Fisher). Cells were subsequently incubated with Annexin V-FITC (Thermo Fisher) for 15 minutes at room temperature in the dark. After staining, cell pellets were resuspended in 1× binding buffer before data acquisition.

Data acquisition for all flow cytometry assays was performed on an LSRFortessa 5L flow cytometer (BD Biosciences) and analyzed using FlowJo v10 (TreeStar).

### Functional in vitro assays

For cytokine release assays, 1×10^5^ tumor cells were seeded in 48-well plates. After overnight incubation, T cells were added at an effector-to-target (E:T) ratio of 3:1. Supernatants were collected 24 h after co-culture and the concentration of human IFNγ and IL-2 was analyzed by ELISA using the following kits: DuoSet ELISA Development kit (R&D Systems, #DY285B and #DY202) following manufacturer’s protocol. Absorbance was determined using Gen5 2.07 (Biotek) or iControl 2.0 (LifeSciences) software in an Infinite 200 PRO Plate Reader (Tecan) set to 450 nm and 570 nm for background subtraction.

The killing activity of CAR-T cells in co-culture with healthy cells was analyzed in vitro using the xCELLigence Real-Time Analyzer System (ACEA Biosciences). For this, 1×10⁴ tumor cells were plated in 50 μL of medium per well of an E-16 plate (Agilent), which had previously been filled with 50 μL of target cell medium to set the blank. After 18 h approximately, effector cells were added in a final volume of 100 μL of CAR-T medium at a 1:1 E:T ratio. For tumor only control, 100 μL of CAR-T medium were added to each well. Cell index was monitored every 20 min for four to six days. Relative cell impedance was normalized to the maximum cell index value before T cell plating.

### qPCR for lentiviral insertion copy number determination

Genomic DNA was extracted from CAR^+^ and CAR^-^ T cell fractions at day 11 of expansion using the DNeasy Blood & Tissue Kit (Qiagen). Following extraction, DNA samples were quantified with a NanoDrop spectrophotometer (Thermo Fisher) and diluted to a concentration ranging from 25 to 100 ng/μl. For each sample, 100 ng genomic DNA was amplified. qPCR was performed using the LightCycler 480 SYBR Green I Master mix (Roche) and custom primers (10 µM, Integrated DNA Technologies). The following primer sets were employed: GATA (Forward (Fw): 5’-TGGCGCACAACTACATGGAA; Reverse (Rv): 5’-CGAGTCGAGGTGATTGAAGAAGA), WPRE (Fw: 5’-CGCTGCTTTAATGCCTTTGTAT; Rv: 5’-GGGCCACAACTCCTCATAAA) and HER2 (Fw: TGGAATGGGTTGCAAGGAT; Rv: TGGATGTGTCTGCGCTTATAG). Standard curves for each gene were generated, ranging from 10^8^ to 10^2^ copies per genome. The reactions were carried out in 384-well plates (Roche) on a LightCycler 480 system (Roche), with cycling conditions consisting of an initial denaturation phase at 95°C for 5 min, followed by 40 cycles of amplification at 95°C for 10 seconds, 58°C for 10 seconds and 72 °C for 5 seconds. The Ct values were calculated using the LightCycler software and normalized to the GATA reference gene. All samples, including those used for standard curves, were analyzed in duplicate. Data acquisition and analysis were conducted using the instrument’s software to determine the cycle threshold (Ct) values for each sample. Ct values were normalized to the reference gene GATA. WPRE was used to quantify the total number of CAR molecules per sample.

### Gene expression analysis

Gene expression analysis was performed using the CAR-T cell characterization panel from Nanostring Technologies (Seattle,WA). Briefly, total RNA was extracted from either control T cells or CAR-T cells at indicated timpoints and either treated or not with dasatinib using the RNeasy Mini kit (Qiagen). Samples were prepared according to the manufacturer’s protocols for the nCounter CAR-T Characterization Panel. Cartridges were run on the nCounter SPRINT Profiler. The raw digital count of expression was exported from nSolver v3.0 software for downstream analysis. Gene expression levels were normalized against the housekeeping genes. Data analysis was conducted in R (v4.3.2). Differentially expressed genes between conditions were explored using the limma package (v3.58.1) incorporating donor as a paired factor in the linear model and using p.value <0.01 as cutoff. Pre-ranked Gene Set Enrichment Analysis (GSEA) was conducted over the ranked list of genes based on the log2 Fold Change obtained from the differential expression analysis, per comparison. GSEA was conducted through fgsea function from fgsea R package (v1.28.0) with default parameters (http://bioconductor.org/packages/fgsea/). GSEA interrogated Gene Ontology database through gage package (v2.52.0). Plots were done in R.

### Mouse xenograft studies

6–8-week-old NOD/SCID/IL2-receptor y chain knockout (NSG) female mice were subcutaneously implanted with 4-5×10^6^ SKOV3 tumor cells in a final volume of 100 μL per flank, in a 50% solution of Matrigel (Corning) in phosphate-buffered saline (PBS). When tumors reached 150–350 mm^3^, mice were treated with a single intravenous injection of 2–5 × 10^6^ control T cells or CAR-T cells in 100 μL of PBS, following previously published protocols [41]. Tumor dimensions were measured weekly with a digital caliper, and volumes were calculated using the formula V = 6 x (L x W2) / π, where L is length and W is width of the tumor. Mice were sacrificed when tumors reached 1500 mm^3^. T cell persistence was analyzed from peripheral blood samples obtained by retro-orbital bleeding at indicated timepoints. Blood samples were mixed with antibodies against T cell markers CD45, CD4 and CD8 in BD Trucount tubes (BD Biosciences, # 340334), and the absolute number of T cells per microliter was calculated following manufacturer’s protocol.

### GMP-grade CAR-T manufacturing

Lentiviral vectors were produced in a cleanroom facility in accordance to GMP guidelines [37]. Apheresis material was processed using the CliniMACS Prodigy® system (Miltenyi), starting with erythrocyte and platelet depletion through density-gradient centrifugation. CD4⁺ and CD8⁺ T cell subsets were then isolated by magnetic bead selection (Miltenyi Biotec, n. 200-070-213 and n. 200-070-215). A minimum of 1 × 10⁸ selected T cells were used to initiate the manufacturing process. Cell cultures were expanded in TexMACS® medium (Miltenyi Biotec n. 170-076-306) supplemented with with IL-7, and IL-15 (Miltenyi Biotec, n. 170-076-184 and 170-076-114). T cells were activated immediately upon culture initiation using TransACT (Miltenyi Biotec, n. 200-076-204) and transduced 24-48 hours later with lentiviral particles. After 48 hours, cells were washed and subsequently maintained under gradually increasing agitation for 7–10 days, until the target cell numbers were achieved. The final CAR-T product was formulated in saline with 2% human serum albumin and cryopreserved with 10% DMSO. In some cases, dasatinib (50 nM) was added on days 3 and 5 during the primary T-cell expansion.

## Statistical analysis

Statistical analysis was performed using GraphPad Prism v9 (GraphPad Sofware Inc.). Data are presented as mean ± S.D. or mean ± s.e.m, as specified. Comparisons between sample means and a defined reference value were performed using a one-sample t-test. Comparisons of multiple groups were performed by using a one-or two-way ANOVA with a Tukey’s multiple-comparisons test. Survival analyses from in vivo experiments were conducted using the Kaplan-Meier method, and statistical differences between groups were assessed with the log-rank test. Specific details regarding the statistical methods are provided in the corresponding figure legends. P-values are represented as follows: *p < 0.05, **p < 0.01, ***p < 0.001 or ****p < 0.0001.

## Data availability statement

The datasets generated and/or analyzed during the present study are available from the corresponding author on reasonable request.

## Acknowledgements

This study was supported by the Spanish Ministry of Science and Innovation under Ramon y Cajal grants RYC2018-024442-I to S.G.; the Spanish Ministry of Health under a Rio Hortega 2020 to L.A, research funding from “la Caixa” Foundation to M.J. (LCF/ PR/SP23/52950004); the Spanish Association Against Cancer (LABAE20022GUED to S.G., INVES222988RODR to A.R.-G. and AECC-IDIBAPS Excellence Program EPAEC246711CLIN to AP), and the AGAUR_INVESTIGO22 100028TC1 grant, within the framework of the Recovery, Transformation, and Resilience Plan, funded by the European Union’s NextGeneration EU Recovery Mechanism. This work was developed at the Centro Esther Koplowitz, Barcelona, Spain. We thank the Flow Cytometry and Cell Sorting core facility of Fundació de Recerca Clínic Barcelona-Institut d’Investigacions Biomèdiques August Pi Sunyer (FRCB-IDIBAPS) for their technical help. We also thank the animal facility of the University of Barcelona.

## Author contributions

L.A. performed experiments, analyzed and interpreted the data, and contributed to writing the original draft of the manuscript. B.M. performed experiments and analyzed and interpreted the data. A.R.-G. analyzed and interpreted the data and wrote the original draft of the manuscript. T.L.-J. performed analysis and visualization of transcriptomic data and contributed to writing the original draft of the manuscript. M.H.-S., G.C., S.C., I.A-S., and M.G.-A. performed *in vitro* experiments. J.C. and G.C. conducted *in vivo* studies. H.C. developed CAR constructs used in the study and produced clinical-grade batches of lentiviral vectors. P.G. helped with NanoString experiments. M.E.-R. and E.A.G.-N. manufactured HER2 CAR-T cell batches under GMP conditions, performed experiments, and analyzed the data. A.U.-I., J.D., A.P., and M.J. provided conceptual guidance. S.G., A.P. and M.J. contributed to funding acquisition. S.G. supervised the project, including conceptualization, experimental design, data analysis, and manuscript writing. All authors revised and approved the final manuscript.

